# The timing network is engaged in the practice of internally driven tapping independently of the learning transfer from perceptual to motor timing

**DOI:** 10.1101/2020.12.17.423301

**Authors:** Itzamná Sánchez-Moncada, Luis Concha, Hugo Merchant

## Abstract

When we intensively train a timing skill, such as learning to play the piano, we do not only produce brain changes associated with task-specific learning, but also improve the performance on other temporal behaviors that depend on these tuned neural resources. Since the neural basis of time learning and generalization are still unknown, we measured the changes in neural activity associated with the transfer of learning from perceptual to motor timing. We found that intense training in an interval discrimination task increased the acuity of time perception in a group of subjects that also showed learning transfer, expressed as a reduction in tapping variability during an internally-driven periodic motor task. However, we also found subjects with no learning and generalization effects, and a third group with no signs of learning but with practice-based decreases in temporal variability in the motor task. Notably, these heterogeneous populations of subjects shared a common increase of activity in the medial premotor areas and the putamen in the post-with respect to the pre-training session of the tapping task. These findings support the idea that the core timing network is constantly refining its ability to time behaviors in different contexts and that practice is critical for keeping the neural clock attuned and properly functioning.

## 1 Introduction

Our brain can flexibly quantify time across complex perceptual and motor behaviors such as the appreciation and execution of music. These behaviors demand the development of sophisticated skills to extract the beat or isochronous pulse of intricate musical patterns and to produce predictive movements entrained to the beat, which can reach exquisite levels of temporal performance in professional percussionists (Honing & Merchant 2014, Mendoza & Merchant 2014). Hence, temporal learning and processing are critical elements of human intelligence that have been investigated for decades (Treisman 1963, Herholz & Zatorre 2012, Ayala et al. 2017). The classical view from experimental psychology of a common clock for timing across sensory and motor tasks (Kristofferson 1980, Ivry & Hazeltine 1995, Gibbon et al. 1997) has been replaced by imaging and neurophysiological studies supporting the notion of a partially distributed neural timing circuit that possesses two elements (Rao et al. 1997, Jantzen et al. 2002, Macar et al. 2006, Coull et al. 2008, Wiener et al. 2010, Merchant, Pérez, Zarco & Gámez 2013). The first element is the core timing network, integrated by key areas of the motor system, namely the cerebellum and the corticothalamic-basal ganglia (CTBG) circuit (Merchant, Grahn, Trainor, Rohrmeier & Fitch 2015, Tanaka et al. 2020). This core timing network is involved in temporal processing in a wide range of perceptual and motor timing behaviors in the hundreds of milliseconds scale, including visual, auditory and tactile stimuli, and a variety of motor effectors (Wiener et al. 2010, Merchant, Harrington & Meck 2013). The second element is represented by areas selectively engaged on the specific behavioral requirement of a task (Buhusi & Meck 2005, Coull et al. 2011, Harrington et al. 2011). These task-dependent areas interact with the core timing system to produce the characteristic pattern of performance variability of a specific timing paradigm (Merchant, Zarco & Prado 2008, Merchant, Harrington & Meck 2013).

The notion of a core timing network has been also supported by experiments that evaluate learning and generalization of timing (Bueti & Buonomano 2014). The hypothesis behind these studies is that the learning-based improvements in temporal processing within a particular task will show transfer to another timing behavior if they share trained neural circuit resources. Normally, learning transfer is quantified as an increase in time precision when comparing temporal performance in the generalization task between a post-versus a pre-training session. This strategy is followed frequently in the artificial neural network literature, namely, after training a recurrent neural network in a condition with specific input-output rules, the network is tested on other conditions to determine generalization capabilities due to common neural weights and shared internal dynamics (Laje et al. 2018, Pérez & Merchant 2018, Bi & Zhou 2020, Merchant & Pérez 2020). Thus, robust temporal generalization, measured from intensive training in time discrimination, has been documented as an increase in timing acuity across auditory frequencies (Wright et al. 1997, Karmarkar & Buonomano 2003), sensory modalities (Nagarajan et al. 1998, Westheimer 1999, Bartolo & Merchant 2009), stimulus locations (Nagarajan et al. 1998), and relevant to the present study from sensory to motor-timing tasks (Meegan et al. 2000, Planetta & Servos 2008, Di Fabio 2011). These findings strongly support the existence of a multimodal and multicontext core timing network (Merchant, Zarco, Bartolo & Prado 2008, Wiener et al. 2010, Merchant & Yarrow 2016).

A critical aspect in learning-generalization protocols and their implementation as intervention procedures is the individual differences in the resulting outcomes, suggesting heterogeneity with at least three groups. A group with preexisting traits that allows them to learn and transfer their learned gain into another task when neural resources are shared between tasks (Learners), a group of subjects that show high initial levels of timing precision that produce ceiling effects in training and generalization (Non-Learners, around one third of subjects (Bueti & Buonomano 2014)), and a third group that show no learning but show a practice based decrease in temporal variability between the two sessions of the generalization paradigm (Covert Rhythmic-Skill Learners, (Grondin & Ulrich 2011)).

Here, we recruited thirty-nine healthy human subjects that underwent intensive interval discrimination training for a week and performed Pre- and Posttraining sessions of a Synchronization-Continuation tapping task inside an MRI scanner. We found a large subpopulation of participants that showed learning gains on the precision of interval discrimination that were transferred to the temporal execution of a motor task with an initial tapping synchronization to a metronome followed by a self-driven rhythmic response. In addition, we also found groups of subjects behaving as Non-Learners and Covert Rhythmic-Skill Learners. Thus, we next focused on the change in hemodynamic responses associated with the transfer of learning from perceptual to motor timing and compare them with the brain activation profiles of the Non-Learner and the Covert Rhythmic-Skill Learner subpopulations in the post versus pre-training sessions.

## 2 Methods

### 2.1 Subjects

41 right-handed healthy subjects (25 women and 16 men), mean age 27 years old (age range: 20-33 years), with no record of neurological or psychiatric disorders, and normal or corrected-to-normal vision, underwent an intensive interval discrimination training for a week and did a Pre- and Post-training Synchronization-Continuation task session inside an MR scanner (Figure 1). Two subjects were excluded from the final analyses because of their high rate of incorrect responses in the perceptual and tapping tasks. All subjects gave written informed consent for the study protocol, which was approved by the bioethics research committee of the Instituto de Neurobiología, UNAM. The study was performed in accordance with the ethical standards of the Declaration of Helsinki.

**Figure 1:**
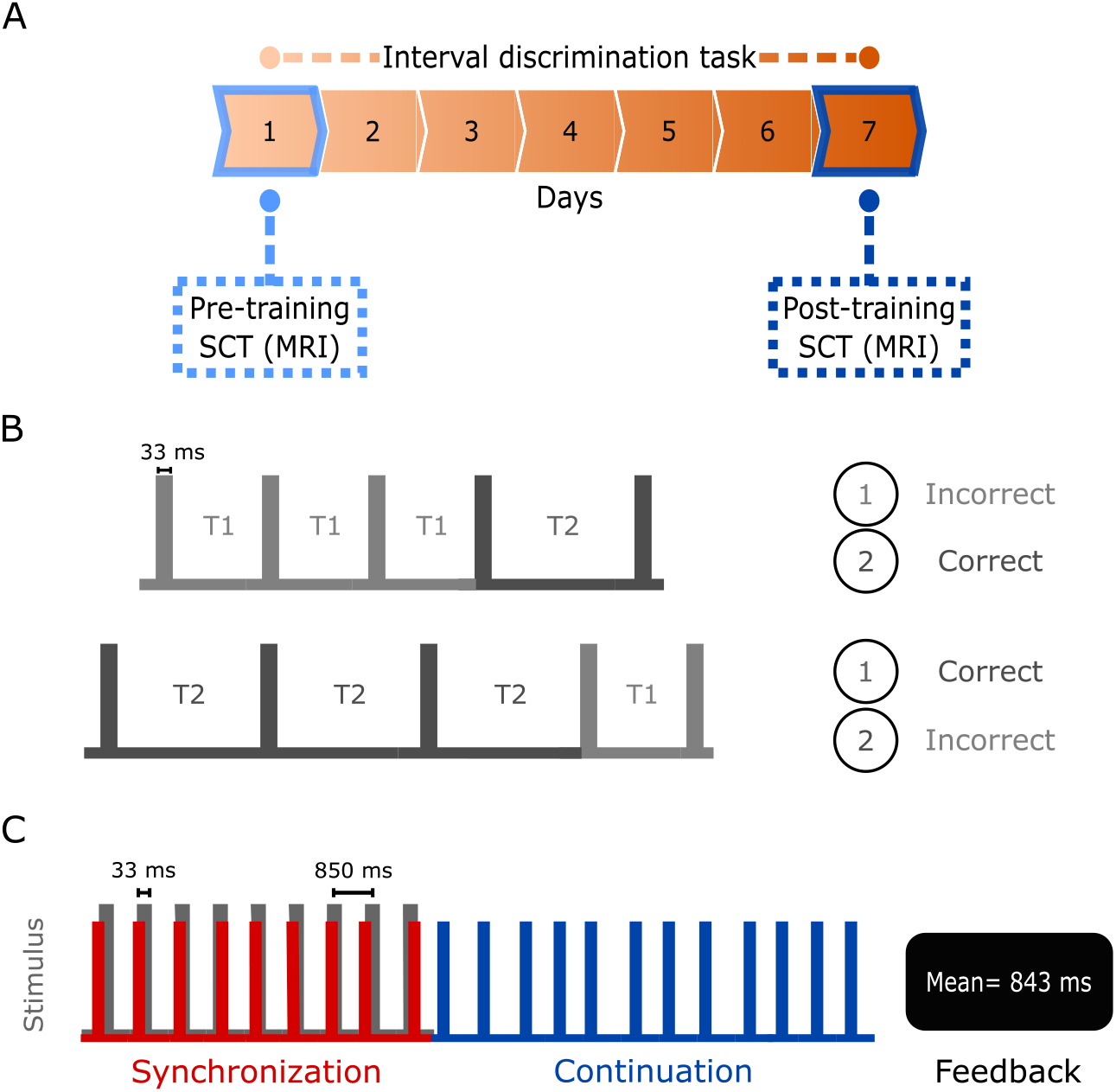
Experimental design. A, Experimental timeline. Pre-training SCT took place on the first day. Later that day, IDT training started, with one-hour daily sessions for 7 days. On the seventh day, Post-training SCT was performed. B, Interval Discrimination Task (IDT). At the beginning of each trial, two different empty intervals (T1 and T2) were presented to the subject. The intervals were delimited by a gray square with a refresh rate of 33 ms. The first interval was presented three times, while the second interval was presented only once. After the last square was shown, the subject answered which one of the two intervals was the longest. C, Synchronization-Continuation Task (SCT). Each trial began with the presentation of a visual metronome consisting of a flashing gray square (33 ms) with a constant inter-stimulus interval of 850 ms. The subject’s objective was to adjust a motor response, pressing a button in synchrony with the visual metronome. After nine taps the visual metronome disappeared and the subject had to continue pressing the button at the same rate, for twelve more times, now internally guided. At the end of each trial in both tasks, feedback was given to the subjects.

### 2.2 Tasks and training

#### 2.2.1 Apparatus

Both tasks were programmed using Matlab R2013a and Psychtoolbox library (Brainard 1997). A Dell XPS Intel Core i5 laptop with Windows 7 was used to run the tasks. During the Interval Discrimination Task, all participants were seated comfortably on a chair facing the laptop with a 15-inch screen in a quiet experimental room; the laptop responding keys were restricted to the space bar, left and right arrow keys.

The Synchronization-Continuation Task was performed inside the MRI scanner. The task was presented through binocular LED screens with diopter correction (VisualSystem; NordicNeuroLab, Bergen, Norway) and responses were registered through a hand-held response collection device (ResponseGrip Nordic-NeuroLab). Subjects were instructed on how to perform the task before entering the scanner and were allowed to do some practice trials with a different ITI interval. Additionally, they were told that the ITI they were performing inside the scanner would be shorter, of 850 ms.

#### 2.2.2 Interval Discrimination Task (IDT)

Subjects discriminated which of two intervals had the longest duration. We employed empty intervals which were delimited by a 3.77 x 3.77 cm2 gray square that flashed at the center of a black screen. The duration on screen of each marker was 33 ms (screen resolution was 1366 x 768 pixels and the refresh rate was 60 Hz). One of the intervals had a constant duration of 850 ms (standard interval), while the other (comparison interval) was selected pseudo-randomly without repetition from the following 8 values: 566, 666, 783, 816, 883, 916, 1033 and 1330 ms. We use the term “repetition” to refer to the subsequent presentation of 8 intervals. Whether the standard interval or the comparison interval was presented first was determined randomly. The first interval was presented three consecutive times, whereas the last interval was presented only once (Figure 1B). To measure the response time, subjects were asked to press and hold the spacebar since the beginning of each trial. Then, subjects had to release the spacebar and press the left or right arrow key to indicate whether the first or second interval was longer, respectively. All actions were performed with the right hand. Feedback was given at the end of each trial on whether the response was correct or incorrect. During a training session, the subjects completed 4 blocks of 10 repetitions (320 total trials, with a duration 60 minutes per training session).

#### 2.2.3 Synchronization-Continuation Task (SCT)

Subjects were lying down inside the scanner with the video goggles comfortably adjusted. At the beginning of each trial, subjects were instructed to fixate an isometric white cross (1.2 cm) that appeared at the center of the black screen. After a variable period (1.2 to 2.4 s) a 3.77 x 3.77 cm2 gray square was presented in sequence as a metronome with an isochronous interstimulus interval of 850 ms. Subjects were instructed to entrain to the visual metronome by pressing a button with their right index finger. After nine synchronized taps (Synchronization epoch) the visual metronome ceased, and the subjects were required to continue pressing the button for another twelve taps (Continuation epoch), attempting to maintain the same beat. The fixation cross was present during both epochs. On the Continuation epoch, once the subject pressed the button for the twelfth time, the fixation cross disappeared, the mean inter-tap-interval (ITI) was calculated and presented to the subject as feedback for 2 s (Figure 1C). Afterwards, the screen was completely black for 10 s (inter-trial interval) and then the white cross appeared again, signaling the start of the next trial. If the asynchronies (the time between the visual cue and the response) were greater than ±425 ms, the trial was excluded from the behavioral and image analysis. Three runs were performed per SCT session, the first one with 20 trials and the rest with 16 trials. Each run lasted for around 10 minutes.

#### 2.2.4 Procedure

A Pre-training/Training/Post-training intervention was implemented (Wright et al. 1997, Bartolo & Merchant 2009). On the first session, subjects performed the SCT (Pre-training) within the MR scanner. Later that day, subjects started their first training session of a training program of seven days on the IDT. On the seventh day and after completing the IDT, subjects performed the second SCT session (Post-training) inside the MR scanner (Figure 1A).

### 2.3 MRI acquisition

Images were acquired in the National Laboratory for Magnetic Resonance Imaging, within our Institution, using a 3.0 T Philips Achieva TX (Best, The Netherlands) system equipped with a 32-channel head coil. A gradient-echo echo-planar imaging sequence (GRE-EPI) was performed to acquire T2*-weighted fMRI images (TR=2 s, TE=30 ms; voxel resolution = 2×2×4 mm3). 32 axial slices comprised each EPI volume. The size of the volume allowed us to scan the entire cerebrum and most of the cerebellum (below the VIIB lobule). 5 dummy volumes were acquired at the beginning of the run for T1 equilibration effects. In addition, a 3-Dimensional-Spoiled Gradient-Recalled Echo (3D-SPGR) sequence was used to obtain high resolution T1-weighted images with a 1 mm3 resolution (TR = 8.15 ms, TE = 3.75 ms; image matrix = 256×256×176), which was used for image registration purposes.

### 2.4 Data Analysis

#### 2.4.1 Behavioral data

##### Interval discrimination task (IDT)

The method of constant stimuli was used to estimate the daily thresholds (Getty 1975). The difference threshold was computed from the psychometric curve, where the probability of long-interval discrimination was plotted as a function of the comparison interval (Merchant, Zarco & Prado 2008, Méndez et al. 2014). A logistic function was fitted to the data and the threshold corresponded to half the subtraction of the interval at 0.75 p and that at 0.25 p (Figure 2), which was computed for each of the four blocks per day. Then, we plotted the threshold across the 7 days of training and fitted a power function *(y = Ax^B^* where y=threshold; A=ten raised to the second polynomial coefficient; x=training days and B=first polynomial coefficient). The learning criteria consisted of a significant P-value (p<0.05; from a F-test) for the adjusted model along with a negative slope which was associated to a reduction of the discrimination threshold, implying an improvement of subject’s ability to discriminate the stimuli.

**Figure 2:**
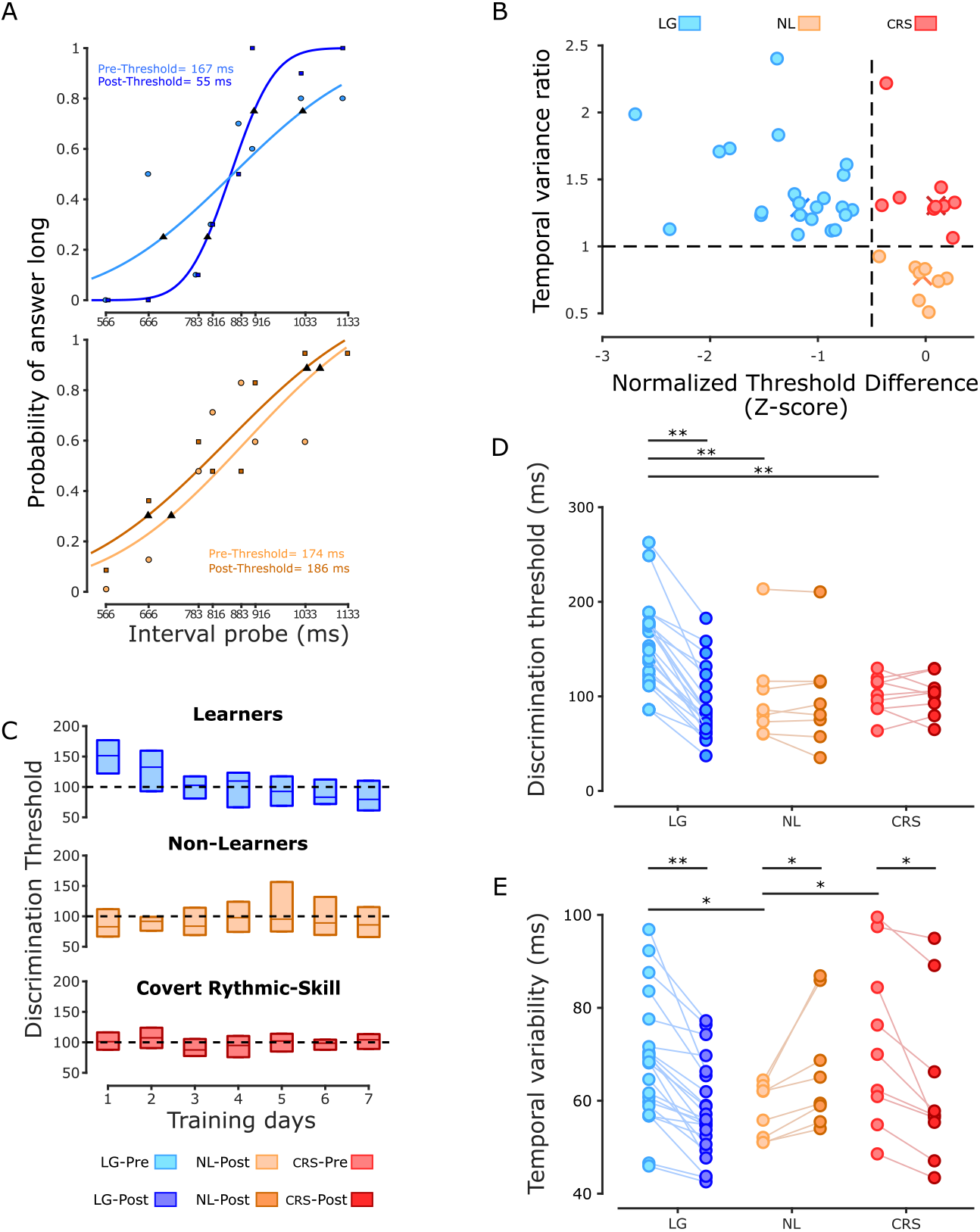
IDT and SCT behavioral analysis. A, Psychometric functions. First and last psychometric function (one block) for a subject with significant reduction of its discrimination threshold (top panel) and a subject without a significant reduction of its discrimination threshold (bottom panel). B, K-means cluster classification identified 3 groups. Blue, orange and red dots correspond to the Learners with Generalization (LG), Non-Learners (NL) and Covert Rhythmic-Skill Learners (CRS), respectively. The horizontal black dotted line corresponds to a Temporal Variance ratio of 1; vertical black dotted line corresponds to a normalized threshold difference of −0.5. Colored X symbols mark the centroid assigned to each group. C, Group discrimination threshold. Interquartile boxplot of the discrimination threshold for subjects which significantly improved their performance for the IDT (Learners group, n=22; top). Interquartile boxplot of the discrimination for subjects without any improvement during the IDT (Non-Learners group, n=8; middle). Interquartile boxplot of the discrimination for subjects without any improvement during the IDT (Covert Rhythmic-Skill Learners group, n=9; bottom). Top and bottom lines of the boxes correspond to the third and first quartile, respectively. D, Discrimination threshold. Discrimination thresholds for each subject, session and group. E, Temporal Variability. Inter-tap variability for each subject, session and group. *p<0.05 and **p≤0.005.

##### Synchronization-Continuation task (SCT)

The first 4 trials of run 1 of SCT were not included in the analysis to obtain data from a steady behavioral response, for a total of 48 analyzed trials (3 runs of 16 trials each). The Synchronization epoch included 9 taps and 8 inter-tap intervals (ITI), but the first tap and ITI were discarded. The Continuation epoch consisted of 12 taps and 12 ITIs, the last ITI was not included in the analysis. In addition, trials were not further analyzed when asynchronies were above ±425 ms (half the duration of the interstimulus interval) or a single ITI was larger than 850 ms ±400 ms. Hence, we got an uneven number of ITIs and taping times per subject. Consequently, a bootstrap resampling method (10,000 iterations) was carried out to get a homogenous number of data points across subjects.

For each subject, we compared the following SCT performance measures between Pre- and Post-sessions: Asynchronies, Constant Error, and Temporal Variability. Asynchronies were the time difference between tap and stimulus onsets and were computed only for the Synchronization epoch. The Constant Error was the average difference between ITIs and the instructed interval. The Temporal Variability was defined as the standard deviation of ITIs. We also computed the Temporal Variance Ratio (TVR) that is the ratio of the ITIs variance of the Pre-divided by the Post-session variance. Therefore, a TVR value below 1 corresponds to an increase, whereas a value above 1 corresponds to a decrease in Temporal Variability in the Post-with respect to the Pre-session. The Constant Error and the Temporal Variability were calculated separately for the Synchronization and Continuation epochs.

Asynchrony values were presented as phases with respect to the beat onset times over the instructed interval. Asynchronies were transformed from milliseconds (*a_i_*) to angular units in radians (*θ_i_*) with the equation *θ_i_* = (2*πa_i_*)/*T_i_*, where *T_i_* corresponded to 850 ms, the target interval. Circular statistics were used to summarize the distribution of the relative phases on the unit circle using the mean resultant vector, which has two parameters the length R (dimensionless ranging from 0 to 1) and angle (given in radians from 0 to 2*π*). R equal to 0 means phases in Asynchronies that are uniformly distributed along the whole inter-onset interval, whereas an R value of 1 indicates identical phases (Figure 3A). A vector angle of 0 meant a perfect temporal alignment between tap and stimulus, while positive and negative angles indicate that the tap followed (positive) or preceded (negative Asynchronies) the stimulus, respectively (Gámez et al. 2018). Asynchronies were analyzed with Matlab’s Circular Statistics Toolbox. The Rayleigh test was used to assess unimodality with the null hypothesis of a uniform distribution around the circle. To assess differences between sessions and groups a Harrison-Kanji test was performed. This test is a parametric two-way ANOVA for circular data with session (Pre- and Post-training) as within-subjects factor and the group as between-subjects factor. One-sample mean angle tests were also performed to determine whether the mean angle of each group was significantly different from zero. Three-way repeated measures ANOVAs were carried out using the Constant Error and the Temporal Variability as dependent variables, the session (Pre- and Post-training) and the epoch (Synchronization and Continuation epochs) as within-subjects factor, and the group (Learners, Non-Learners and Covert Rhythmic-Skill Learners, see below) as between-subjects factor. Two-way repeated measures ANOVAs were performed using the Constant Error and the Temporal Variability as dependent variables, separately, the session (Pre- and Post-training) as within-subjects factor and the subjects*’* group (Learners, Non-Learners and Covert Rhythmic-Skill Learners, see below) as between-subjects factor for the Synchronization and Continuation epochs, separately (Figure 3B and C). Paired t-tests were used as post-hoc test to assess the differences between groups and sessions. Routines for statistical analysis were written using Matlab R2013a. The statistical level to reject the null hypothesis was *α*=0.05.

**Figure 3:**
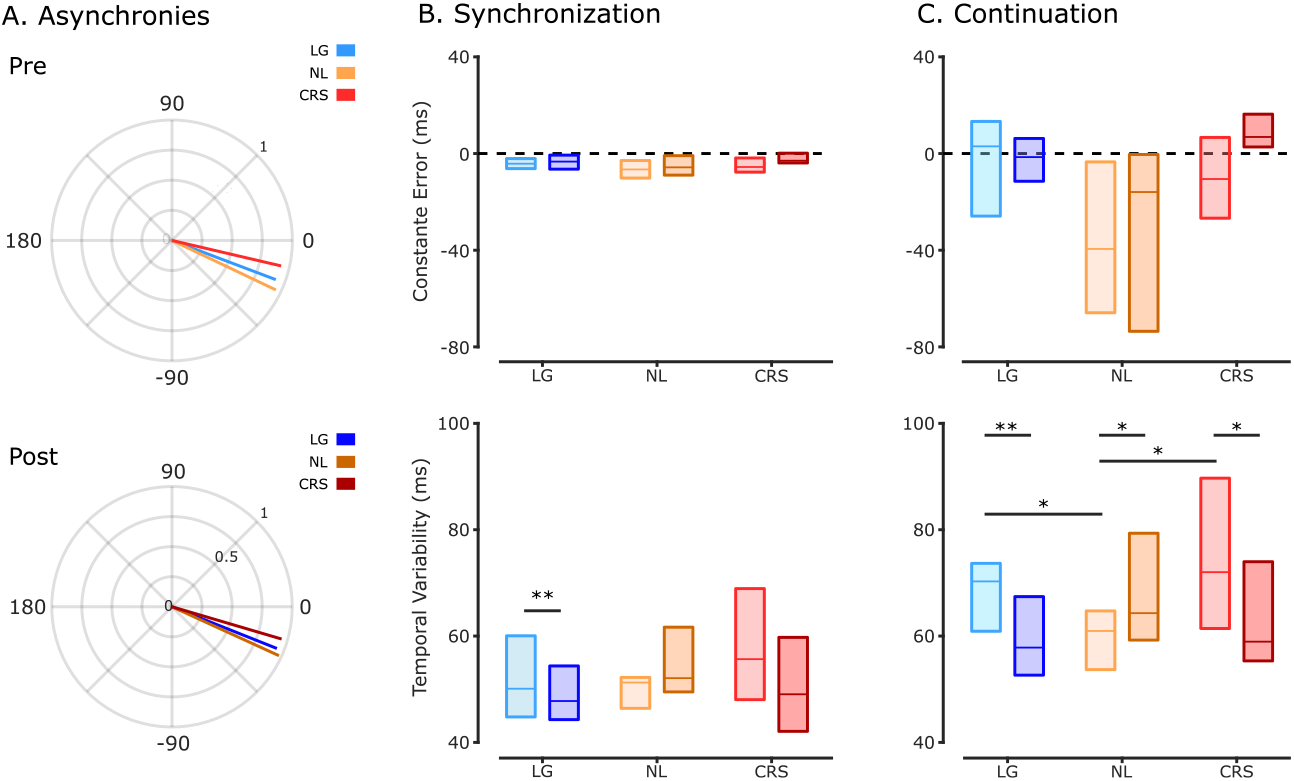
SCT behavioral analysis. A, Asynchronies. Synchronization epoch Asynchronies for the three groups and for the Pre- (top) and Post- (bottom) training sessions. B, Synchronization epoch. Constant Error calculated for all three groups during the Pre- and the Post-training sessions (top). Temporal Variability (standard deviation) for each of the three groups during the Pre- and Posttraining sessions (bottom). C, Continuation epoch. Constant error calculated for the three groups and both sessions (top). Temporal Variability calculated for the three groups and both sessions (bottom). *p<0.05 and **p≤0.0005.

### 2.5 Behavioral clustering

We plotted the temporal variance ratio (TVR) of the Continuation epoch of the SCT as a function of the normalized threshold difference (z-score) between the first and last days of training in the IDT. The former is a measure of temporal generalization with values above 1 indicating a decrease in Temporal Variability in the Post-with respect to the Pre-training session. The latter is a measure of temporal learning, with values below 0 indicating an increase in temporal acuity as a result of a decrease in the discrimination threshold after the daily intensive training. We found the two expected groups of subjects based on previous studies (Meegan et al. 2000, Planetta & Servos 2008). First, we identified a group of Learners with a negative threshold difference statistically different from zero (see also the above Learner criteria) with a concomitant time generalization effect, where the TVR was larger than 1 and a significant effect of session (permutation test). This group was called Learners with Generalization (LG, n=22, blue dots in Figure 2B). Second, a group of subjects with no learning, with a threshold difference that was not statistically different from zero, and no time generalization (TVR below 1), named Non-Learners (NL, n=8, orange dots in Figure 2B). Notably, we also found a third group that were Non-Learners with a TVR with a significant decrease in the post training session, and therefore called Covert Rhythmic-Skill Learners (CRS, n=9, red dots in Figure 2B). We ran a *k*-means clustering with a ‘City Block distance’ metric and *k*=3. The Centroids in *x* and *y* coordinates (normalized threshold difference and TVR, respectively) were: centroid 1 = [-1.1, 1.2], centroid 2 = [0, 0.7], centroid 3 = [0, 1.3] (Figure 2B).

### 2.6 fMRI data analysis

#### 2.6.1 Pre-processing

Pre-training and Post-training functional imaging data were analyzed using Oxford Centre for Functional MRI of the Brain Software Library v5.0 (FSL). All EPI volumes were time and motion corrected. All images were resampled to 2-mm isotropic voxel size and were spatially smoothed using an isotropic Gaussian kernel of 6 mm full-width half-maximum (FWHM) to increase their signal-to-noise ratio.

#### 2.6.2 First-level analysis

An event-related analysis was carried out. Three regressors were used to model the Synchronization, Continuation, and Feedback epochs. All tapping responses were modelled as one event of the correspondent epoch. Each regressor was convolved with a double-gamma function that accounted for the hemodynamic response function. A low-frequency filter was adjusted to the data for any physiological drift (high-pass filter of 100 seconds). Nuisance regressors included 24 motion parameters, estimated by MCFLIRT motion correction. Statistical parametric maps derived from the general linear model were created for each subject during task performance. T-statistics were calculated and then transformed to Z-score maps. First-level analysis was run for each of the 39 subjects to define patterns of activation as compared to baseline.

#### 2.6.3 Second-level analysis

Contrast parameter estimate (COPE) maps of the first-level analysis were averaged for each session and subject across the three SCT runs. These COPEs were utilized to perform the rest of the second-level analysis (unless indicated otherwise). As an initial step, the mean group activation of all subjects was calculated for the Synchronization and the Continuation epochs, separately, only for the Pre-training session. The goal was to identify the areas involved in the two epochs of the SCT before IDT training (Figure 4). A multiple comparisons correction was implemented via AFNI’s 3dttest++ -clustsim tool to estimate the minimum cluster size by Monte-Carlo simulation, with a threshold Z-score of 3.7 (p=0.0001) and a family-wise error rate (FWE) of 0.05 (Eklund et al. 2016, Cox et al. 2017). The critical cluster size was 26 voxels (208 mm^3^) for both Synchronization and Continuation activation maps (Figure 4).

**Figure 4:**
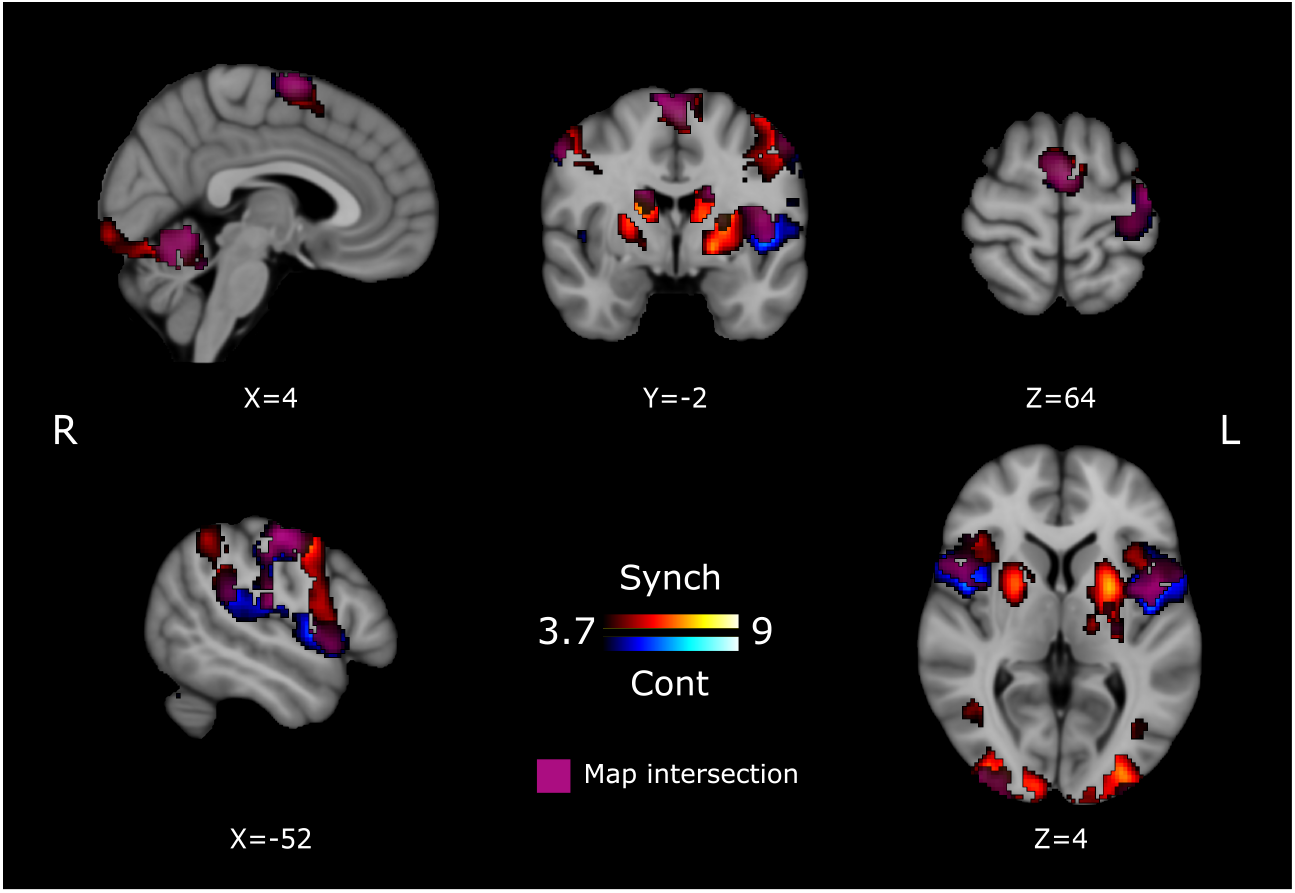
Pre-training group mean activation for Synchronization and Continuation epochs. Warm colors represent areas that were active during the Synchronization epoch. Cool colors represent the areas that were active during the Continuation epoch. Purple denotes areas that were commonly activated for both epochs. Activation maps are displayed as Z-scores thresholded at the cluster-level, overlaid on the MNI template.

A one-way ANOVA was performed between the three behavioral groups for the Pre-training session during the Continuation epoch. The goal was to identify areas that could explain the behavioral differences before any experimental manipulation. Additionally, the COPEs from the first-level analysis were entered for a second-level analysis paired t-test contrasting the Pre-against the Post-training activity for each subject. With the COPEs from the contrast Post > Pre, a third-level one-way ANOVA was performed to assess the differences between groups (LG, NL and CRS) after a week of training. Finally, with the 39 subjects group we performed a two-sample paired t-test to determine the areas that significantly changed their activity between the Pre- and the Post-training sessions during the Synchronization and the Continuation epochs, separately. Multiple comparison correction threshold Z-score was 2.57 and an FWE of 0.05 (Figure 5). Only the Post > Pre contrast during the Continuation epoch survived the multiple comparison correction with a cluster size of 721 (5768 mm^3^).

**Figure 5:**
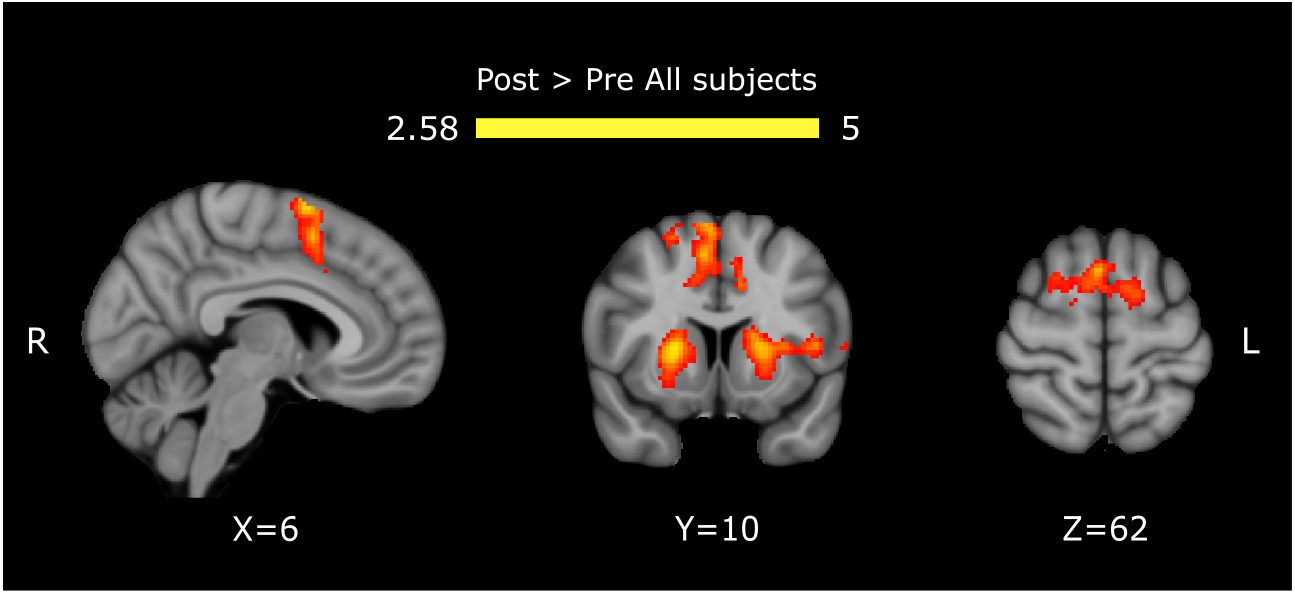
Functional changes related to the SCT. Post-training > Pre-training activity in the Continuation epoch. Areas with increased activity after training all subjects. Activation maps are displayed as Z-scores thresholded at the cluster-level, overlaid on the MNI template.

## 3 Results

### 3.1 Behavioral data

The first goal of this study was to determine whether intensive practice improved interval discrimination performance. Figure 2A (top and bottom panels) depicts the psychometric functions of two participants during the first and the last days of training. While some subjects showed an increasing psychometric function slope as training progressed (Figure 2A, top panel), consistent with a decreased of the discrimination threshold due to training, other subjects failed to show this progressive increase (Figure 2A, bottom panel). Two general groups of subjects were observed: The Learners group (n=22) with subjects that fulfilled the learning criteria (see Methods), and the Non-Learners group (n=17) who did not meet them. These results support the notion that intense interval discrimination training in a group of subjects could improve the inner representation of the trained interval which makes easier to discriminate between the standard and the comparison interval.

The next step was to determine whether improved performance in the time perception task could be accompanied by a gain in the SCT. The hypothesis was that Learners should decrease their ITI’s Temporal Variability during the Post-training session of the SCT, whereas Non-Learners should show similar Temporal Variability between the Post-and Pre-training sessions. Consequently, in Figure 2B we plotted the normalized difference in the discrimination threshold between the last and first days of IDT training against the Temporal Variance Ratio (TVR) of the Continuation epoch of the SCT. The TVR is a measure of temporal generalization (the ratio Pre-/ Post-training ITI variance) with values above 1 indicating a decrease in Temporal Variability in the Post-with respect to the Pre-training. Complex inter-subject differences are evident in Figure 2B and an iterative k-means clustering determined the following three groups of subjects: 1) 22 subjects who reduced their discrimination threshold and their ITI variability, called Learners with Generalization (LG); 2) 8 subjects who did not reduce their discrimination threshold nor their ITI variability, called Non-Learners (NL) and 3) 9 subjects who did not reduce their discrimination threshold but were able to reduce their ITI variability, called Covert Rhythmic-Skill Learners (CRS).

After the three groups were defined, we found statistically significant differences between the groups during the IDT. A two-way ANOVA on the interval discrimination threshold showed significant main effects for session (*F*_1,36_=31.786, p<0.0001) and for session x group interaction (*F*_2,36_=39.388, p=<0.0001) (Figure 2D). Post-hoc paired t-tests showed a significant threshold reduction for the LG (*t*_21_=11.2707, p<0.0001). No significant changes were found for NL (*t*_7_=0.5375, p=0.6076) and CRS (*t*_8_ =-0.0223, p=0.9827) group. Additionally, Pre-training discrimination thresholds were significantly higher for the LG compared to the ones of the NL (*t*_28_=2.9346, p=0.0066) and CRS (*t*_29_=3.451, p=0.0017), but not between NL and CRS (*t*_15_=-0.1068, p=0.9164). Hence, these results confirm the existence of a large group of Learners (LG) with higher initial discrimination threshold that is reduced after a week of intense interval discrimination training. On the other hand, the group of NL and CRS started training with significant lower initial discrimination threshold, high discriminant capabilities, that could account for their inability to increase their performance after training (ceiling effect).

Asynchronies correspond to the time difference between stimulus onset and tap onset during the Synchronization epoch across groups and sessions, as serve a measure of sensorimotor prediction. The mean Asynchronies were plotted as relative phases on the unit circle across Groups and Sessions (Figure 3A). We found that the mean resultant was close to one indicating a consistent synchronization to the metronome among all groups (Rayleigh test for the Pre-training Asynchronies: z=5096, p<0.0001 (LG); z=1990, p<0.0001 (NL); z=2126, p<0.0001 (CRS). Post-training Asynchronies: z=5290, p<0.0001 (LG); z=2088, p<0.0001 (NL); z=2216, p<0.0001 (CRS)). In addition, the three groups in the Pre- and Post-Training sessions showed negative mean circular Asynchronies (One-sample mean angle test was significantly different from 0 for all groups and sessions, p < 0.05), reflecting a strong predictive behavior in all subjects in the two SCT sessions. Finally, a Harrison-Kanji test (two-way ANOVA for circular data) on the Asynchronies showed no significant main effect for group (*F*_1,72_=0.119, p=0.7311) nor for session (*F*_2,72_=1.0907, p=0.3415). These findings suggest that the predictive mechanisms behind consistent and negative mean Asynchronies are not influenced by generalization of a time discrimination task.

Next, we found that the tapping accuracy is not affected by intense interval discrimination training. A three-way ANOVA, with Constant Error (the difference between produced and 850 ms instructed interval) as dependent variable, showed a significant main effect for group (*F*_2,144_=8.31, p=0.0004), epoch (*F*_1,144_=7.21, p=0.0081), and a significant group x epoch interaction (*F*_2,144_=5.77, p=0.0039). These results support the notion of an accurate estimation of the interval during the Synchronization epoch across groups and sessions, accompanied by a decrease in Constant Error in the Continuation, especially for NL, but without session effects (Figure 3B and C, top). Hence, the interval discrimination learning did not generalize as changes in tapping accuracy across groups of subjects nor SCT epochs.

Regarding the Temporal Variability (standard deviation of the produced intervals), results showed that interval training differentially modifies the tapping precision of the subjects during the Continuation epoch of the SCT. A three-way ANOVA, with the Temporal Variability as dependent variable showed significant main effects of epoch (*F*_1,144_=26.92, p<0.0001), and a significant group x session interaction (*F*_2,144_=5.12, p=0.0071). These effects were mainly due to an increase in Temporal Variability in the Continuation epoch, with heterogeneous changes by session for the different groups, but mainly during internally timed tapping. Consequently, we focused on the changes in the precision of produced intervals between the Pre- and Post-training across groups for this SCT epoch. The corresponding two-way ANOVA showed significant main effects for session (*F*_1,36_=7.116, p=0.0113) and for the session x group interaction (*F*_2,36_ =20.005, p<0.0001). Post hoc paired t-tests showed a significant reduction of the ITI standard deviation between the Pre- and Post-training sessions of LG (*t*_21_=6.6634, p<0.0001), and CRS (*t*_8_=3.6519, p=0.0065), while the NL group showed a statistically significant increase between sessions (*t*_7_= – 3.0174, p=0.0195) (See Figure 3C bottom). These results uphold the notion that intense interval discrimination training is one of the mechanisms that can effectively reduce Time Variability during the internal driven epoch of the motor tapping task. The fact that CRS’ Time Variability decreased without any discrimination threshold reduction may stand for an unmeasured learning phenomenon that is driving the Continuation epoch behavioral improvement. Moreover, initial inter-tap variability of the LG was greater compared to the NL (*t*_28_=2.0641, p=0.0484) but not to the CRS (*t*_29_=-0.8124, p=0.4232). On the other hand, CRS’ inter-tap variability was also greater than the NL’s (*t*_15_=2.2197, p=0.0423). This outcome reveals that LG and CRS’ Time Variability starting point allowed them to increase their precision while performing the Continuation epoch and may be the cause of NL’s incapability to increase theirs.

Overall, these results underpin that the gain in time precision due to intensive interval discrimination training could be transferred as a reduction in Temporal Variability of produced intervals during a motor task. Notably, this learning transfer is limited to the Continuation epoch, where subjects internally produced a sequence of taps with a regular tempo. Important individual differences were observed, with subjects learning during IDT and generalizing during the Continuation of SCT, non-learning IDT subjects with no changes in SCT, and a group of Non-Learners that showed procedural changes in the SCT. Next, we characterized the modulation in BOLD activity between sessions across these three groups.

### 3.2 fMRI data

Our first approach on the functional imaging data was to determine the brain areas involved in the Synchronization and Continuation of taps before training in the interval discrimination task, using a whole brain analysis. All subjects were grouped, and the mean activation was calculated for the Synchronization and Continuation epochs. For both task epochs, areas that showed a statistically larger activation with respect to rest condition included the bilateral SMA, bilateral pre-SMA, left M1, left S1, bilateral dPMC, bilateral vPMC, left planum temporale, bilateral caudate, bilateral BA44 (Broca’s area), bilateral insula, bilateral visual cortices, bilateral cerebellar lobules I-VI and the left Crus I (Figure 4, Table 1 and 2). Although the statistical maps for both task epochs had almost the same pattern of activation, areas of activation were slightly larger in the Synchronization map. Moreover, activation of the basal ganglia (putamen, globus pallidus, motor thalamus), was bilateral in the Synchronization epoch, but was limited to the left hemisphere during the Continuation epoch (Figure 4). These results support the idea that the execution of motor timing tasks rely on a cortico-basal ganglia circuit as well as a cerebellar circuit which are key elements of the core timing system.

**Table 1:** Synchronization activation.

| Volume (mm<sup>3</sup>) | Region | $x$ | $y$ | $z$ | Z-score |
| --- | --- | --- | --- | --- | --- |
| 47,232 | C VI R | 24 | -54 | -24 | 8.27 |
|  | C VI L | -28 | -66 | -22 | 7.54 |
|  | V4 L | -32 | -84 | -2 | 6.95 |
|  | V1 R | 26 | -90 | 4 | 6.65 |
| 41,200 | M1 L | -40 | -20 | 52 | 7.80 |
|  | Put L | -24 | 0 | 12 | 7.50 |
|  | Caud L | -14 | -8 | 20 | 7.35 |
|  | Ins L | -40 | -4 | 18 | 6.89 |
|  | dPMC L | -42 | -8 | 58 | 6.84 |
| 19,992 | Caud R | 18 | 0 | 16 | 7.12 |
|  | Put R | 22 | 4 | 8 | 6.55 |
|  | vPMC R | 54 | 0 | 42 | 6.27 |
|  | BA44 R | 48 | 16 | -2 | 5.51 |
| 6,856 | SMA L | -2 | 0 | 62 | 8.01 |
| 4,896 | IPL L | -42 | -42 | 26 | 5.43 |
| 760 | IPL R | 52 | -40 | 48 | 4.49 |
| 456 | PT R | 62 | -34 | 20 | 4.97 |
| 224 | SPL L | -22 | -54 | 48 | 4.24 |

**Table 2:** Continuation activation.

| Volume (mm<sup>3</sup>) | Region | $x$ | $y$ | $z$ | Z-score |
| --- | --- | --- | --- | --- | --- |
| 37,904 | M1 L | -38 | -20 | 52 | 7.42 |
|  | Ins L | -46 | 4 | -2 | 6.92 |
|  | S2 L | -56 | -16 | 18 | 6.62 |
| 9,896 | C V R | 10 | -52 | -18 | 7.25 |
|  | C VI R | 20 | -56 | -22 | 6.86 |
|  | C I-IV R | 6 | -46 | -26 | 5.19 |
| 8,848 | BA44 R | 50 | 12 | 6 | 6.37 |
| 4,816 | pre-SMA L | 2 | 2 | 68 | 6.45 |
|  | SMA R | 6 | -2 | 68 | 6.45 |
|  | SMA L | -2 | -4 | 64 | 6.43 |
| 3,896 | C VI L | -28 | -66 | -22 | 5.68 |
|  | V2 L | -28 | -98 | -8 | 4.76 |
|  | V3 L | -30 | -100 | 0 | 4.71 |
| 3,008 | PT R | 58 | -32 | 22 | 5.87 |
| 2,840 | V3 R | 40 | -92 | 2 | 5.51 |
|  | V2 R | 30 | -100 | 2 | 5.19 |
| 2,024 | Caud R | 16 | -10 | 24 | 6.00 |
| 1,776 | Caud L | -18 | -6 | 24 | 6.74 |
| 1,072 | Put L | -22 | -2 | 10 | 4.48 |
|  | GP L | -22 | -10 | 2 | 4.39 |
| 1,064 | vPMC R | 56 | -2 | 42 | 5.73 |

We searched for intrinsic differences between LG, NL and CRS groups, with a one-way ANOVA between the three groups for the Pre-training session during the two epochs of the SCT. No significant differences were found between groups for either the Synchronization or the Continuation epochs. These findings indicate that no functional inherent activity could explain the behavioral performance differences between groups, given the experimental conditions and the proposed methodology.

Next, we looked for differences between the three groups for the contrast Post > Pre during the Synchronization and Continuation epochs with a oneway ANOVA. No differences were found between groups. Although behavioral improvements were found in the LG and CRS groups, these changes were not followed by any measurable dissimilarity in the BOLD signal between groups.

Finally, a third-level analysis for all-subjects for the contrast Post > Pre on the Continuation epoch showed significantly increased BOLD signal activity after training. We found increased activity in areas correspondent to the motor cortical and subcortical areas (bilateral preSMA, right pre-motor cortex, vPMC, left paracingulate gyrus, bilateral putamen, left insula and left BA44) (Figure 5, Table 3). These results suggest a larger engagement of main structures of the core timing system during the Post-training session. Furthermore, the increased activity is independent of the learning and time generalization processes and may suggest functional changes only related to operational learning linked to the two practice sessions of the SCT in all the subjects.

**Table 3:**
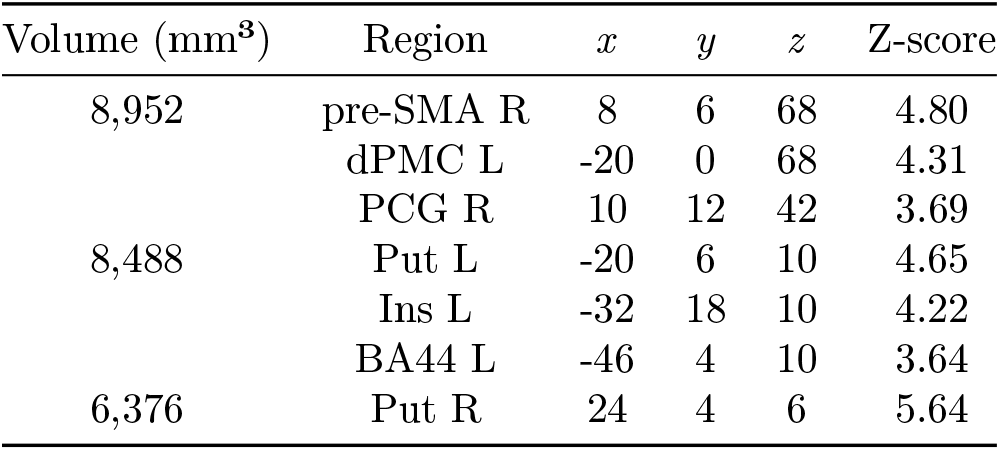
All subject’s contrast Post>Pre.

| Volume (mm<sup>3</sup>) | Region | $x$ | $y$ | $z$ | Z-score |
| --- | --- | --- | --- | --- | --- |
| 8,952 | pre-SMA R | 8 | 6 | 68 | 4.80 |
|  | dPMC L | -20 | 0 | 68 | 4.31 |
|  | PCG R | 10 | 12 | 42 | 3.69 |
| 8,488 | Put L | -20 | 6 | 10 | 4.65 |
|  | Ins L | -32 | 18 | 10 | 4.22 |
|  | BA44 L | -46 | 4 | 10 | 3.64 |
| 6,376 | Put R | 24 | 4 | 6 | 5.64 |

## 4 Discussion

The present research examined changes in neural activity associated with the transfer of learning from perceptual to motor timing and compared with the hemodynamic response of Non-Learners and the Covert Rhythmic-Skill Learners. Our study supports four conclusions. First, intense training in an interval discrimination task produced an increase in the acuity of time perception in a group of subjects considered Learners. Second, there is a strong correspondence between the reduction of the discrimination threshold in the IDT and the reduction of temporal variability of produced intervals during the internally driven epoch of the SCT. Third, initial interval discrimination performance accounted for the lack of learning in Non-Learners and Covert Rhythmic-Skill Learners, evidencing a floor effect on time perception. Last, functional changes occurred in two key areas of the core timing network, the bilateral preSMA and bilateral putamen, were linked to practice effects due to executing twice the SCT in the three groups of subjects and not to the learning transfer of time precision.

Our psychophysical results revealed that a learning process occurred during the seven consecutive days of intensive training in the interval discrimination task. Notably, the learning function of the present study is similar to the time course of learning for auditory, visual, and somatosensory interval discrimination (Kristofferson 1980, Wright et al. 1997, Nagarajan et al. 1998, Westheimer 1999, Karmarkar & Buonomano 2003). All these experiments included intensive daily training for five or more days. With this protocol, learning is characterized by an increase in time perception acuity and occurred mainly during an initial rapid-improvement stage that lasted for 2 or 3 days, followed by a slower improvement phase that spanned the remaining sessions. Nevertheless, important individual differences are also evident in these studies, with a proportion of participants showing no ability to learn and decrease their temporal precision during time perception training. In our case, the Learners and Non-Learners were testing groups that allowed us to investigate not only the generalization rules of timing from perception to production, but also to contrast the neural circuits involved in learning transfer versus those involved in the SCT practice during the Pre- and Post-training sessions.

The SCT has been a prototypical paradigm that contains an initial tapping Synchronization epoch, where subjects entrained to an isochronous metronome, followed by an internally driven Continuation epoch (Wing, 2002; Repp, 2005). Thus, a natural question is whether learning generalization from time perception was present in either or both SCT epochs. The performance in this tapping task can be characterized in terms of precision (Temporal Variability), accuracy (Constant Error), and predictability (Asynchronies specific of the Synchronization epoch) (Zarco et al. 2009, Gámez et al. 2018, Yc et al. 2019). Importantly, the performance gain in temporal precision in our visual IDT was only trans-ferred as an increase in timing precision during the internally driven period of the SCT, but no changes in timing accuracy during this epoch. In addition, no generalization was observed for the precision, accuracy, or predictability during the Synchronization epoch. Furthermore, the increase in timing precision of the Continuation epoch due to learning transfer is evident in the Learner but not the Non-Learner group. Therefore, these results revealed a very specific mechanism for time generalization in Learners: intensive training produced a more robust neural representation of an interval and a concomitant increase in perceptual acuity for this duration. In turn, the improved neural representation of the interval is transferred as an increase in temporal precision when subjects access this neural signal to produce internally driven rhythmic movements. The main question, then, is how this could be achieved? The learning-generalization literature concurs in the principle of lack of generalization in the time domain, where the learned gain in temporal precision does not transfer for durations differing for more than 50% of the trained interval. Indeed, in a previous study we found that training in an interval reproduction task produced a Gaussian generalization function, with large generalization for closely neighboring untrained intervals and no generalization for intervals very distant from the trained duration (Bartolo & Merchant 2009). Therefore, these observations suggest the existence of neural circuits that are tuned to specific time length. In fact, interval tuned cells have been recorded in pre-SMA/SMA (Mita et al. 2009, Merchant, Pérez, Zarco & Gámez 2013, Crowe et al. 2014, Gámez et al. 2019), putamen (Bartolo et al. 2014), caudate, and cerebellum (Kunimatsu et al. n.d.). Thus, the increase in timing precision of an interval during learning and generalization may depend on an increase in the density of neurons tuned to this interval, a decrease in the width of the tuning function of these cells, and/or a concomitant change in the precision of neural population signals across areas of core timing circuit (Merchant, Pérez, Zarco & Gámez 2013, Merchant et al. 2014, Merchant, Pérez, Bartolo, Méndez, Mendoza, Gámez, Yc & Prado 2015, Sohn et al. 2019).

Many functional imaging studies have shown an activation of the core timing circuit during both epochs of the SCT (Rao et al. 1997, Jäncke et al. 2000, Jantzen et al. 2004, Lewis et al. 2004, Gompf et al. 2017). Congruent with these reports, we observed an increase in hemodynamic response during Synchronization and Continuation across SMA/pre-SMA, basal ganglia, ventral and dorsal portions of premotor cortex, inferior parietal cortex and large portions of the cerebellum. Notably, auditory areas were also active during SCT, despite our use of a visual metronome. All these areas on the core timing network were engaged in the SCT execution across the three subject groups, indicating no preexisting traits in this network that distinguish the Learners from the NonLearners and the Covert Rhythmic-Skill Learners. Activation of the superior temporal gyrus (STG), found in both epochs of the SCT, is in line with previous reports on the participation of this area in the performance of auditory and visual rhythms (Rao et al. 1997, Bengtsson et al. 2005). It has been proposed that the STG, a node in the dorsal auditory pathway, could be translating visual presented rhythms into auditory-motor representations (Karabanov et al. 2009). In addition, visual areas were not only active during the sensory cued epoch but also during the internally driven part of the task. These findings support the notion that rhythmic entrainment depends on the active interplay between dorsal auditory stream and the premotor system generating a dynamic bottom-up and top-down system for beat perception and entrainment (Patel & Iversen 2014, Merchant & Honing 2014, Merchant, Grahn, Trainor, Rohrmeier & Fitch 2015). The current hypothesis is that internal predictive timing exists within the motor system, specifically in the CTBGc that constitutes the core timing network (Wiener et al. 2010, Coull et al. 2011). Neurophysiological observations in behaving monkeys indicate that this core timing network encodes an isochronous beat and controls rhythmic tapping throughout neural population dynamics (Merchant & Averbeck 2017). The activity of SMA populations during SCT form periodic state trajectories that behave as regenerating tangent circles, which converge on an attractor that predicts the time of the synchronized tap to each beat event, while changing in amplitude and not speed to represent the tempo (Gámez et al. 2019). On the other hand, the dorsal auditory stream calibrates the core timing system when the internal prediction does not match the rhythmic input. Thus, changes in phase or tempo of the input metronome can be sensed by auditory areas, providing an error signal that is used to flexibly adjust the internal prediction of the beat by the CTBGc and correct the rhythmic output behavior (Morillon et al. 2014). The present results also highlight the privileged position of the auditory system on temporal processing (van Wassenhove & Nagarajan 2007, Kanai et al. 2011) and its engagement in beat-based timing and the SCT, even when we used a visual instead of an auditory metronome, as formerly reported (Karabanov et al. 2009). Finally, the observed activation of the visual areas suggest that the visual system receives a strong top-down predictive signal not only during Synchronization but also during the fully internal control of rhythmic tapping.

In the present study we did not find specific increases or decreases in brain activation profiles for Learners. The three groups of subjects showed similar changes in the BOLD signal of bilateral SMA and bilateral putamen between the post and the pre-learning sessions. At face value, these results suggest that the learning transfer from time discrimination to the internally driven epoch of the tapping task was not accompanied by characteristic changes in the hemodynamic response. This could be due to at least three reasons. First, if we assume as correct the hypothesis (delineated above) that intensive training produces a robust neural representation of an interval, which in turn generates more precise timing in the Continuation of the SCT, it is quite possible that the changes in the response amplitude or the width of interval tuned neurons of the core timing network cannot be detected by the BOLD signal (Logothetis et al. 2001). Ideally, high density single cell recordings in SMA/preSMA and the other areas of the CTBGc (Mendoza et al. 2016) during learning-generalization protocols could provide the data to accept or reject this hypothesis. However, this can only be accomplished in patient populations undergoing neurological surgery or in animal models, with concomitant behavioral and technical problems. Second, a recent elegant study showed that sequence motor learning in human subjects is not associated with increases in BOLD signal (Berlot et al. 2020). Instead, the fMRI correlate of training-induced plasticity corresponds to subtle changes in activity patterns of the cortical premotor system after weeks of digit sequence practice (Berlot et al. 2020). Under this scenario, intensive training in the time discrimination task could produce changes in voxel-to-voxel patterns of activation in areas of the CTBGc, and these plastic changes in pattern activation could be also profited during the Continuation of the SCT to produce the transferred increase in timing. In this line of thought, it is worth mentioning that many imaging studies have found that neural activity decreases with training. Evidence from PET (Jenkins et al. 1994, Jueptner et al. 1997), fNIRS (Ikegami & Taga 2008, Ono et al. 2015), task-related fMRI (Toni et al. 1998, Jantzen et al. 2002, Doyon et al. 2003) and resting state fMRI (Sun et al. 2007, Tamás Kincses et al. 2008, Ma et al. 2011) have found reduction of brain activity after a motor-learning intervention. The fact that we were unable to measure any statistically significant differential activation between groups may be due to an already consolidated learning process (regarding Learners) whose activation profile was not different from the activity detected in Non-Learners nor from the Pre-training session activity. Finally, the training effect of executing the SCT on the two sessions produce a core timing network activation across subject subgroups that may be larger than any specific profile of activation associated with the process of learning transfer from a perceptual to a motor timing task. Indeed, when comparing the hemodynamic response only for Learners we found larger activity between the Post-versus the Pre-training session in the cerebellum and the visual cortex. It should be noted that the cerebellar functional changes we observed were located on the superior cerebellum. Superior and medial cerebellum has been linked to time processing functions during perceptual and motor tasks, mainly in visual modality (Penhune et al. 1998, Jäncke et al. 2000, Schubotz et al. 2000, Lewis & Miall 2003, Lewis et al. 2004, Bengtsson et al. 2005). Lesion (Harrington et al. 2004, Gooch et al. 2010, Schwartze et al. 2016) and transcranial magnetic stimulation (Théoret et al. 2001) studies have shown that medial and superior parts of the cerebellum are implicated in the representation of time. Moreover, non-human primate retrograde tracer experiments revealed a connection between the superior cerebellum (lobules V and VI) and the medial pontine nuclei, which in turn receive their input from occipital areas (Schmahmann 1996). These findings support the notion that the cerebellum, as part of the core timing network, is responsible for encoding a precise representation of the trained duration. In turn, this improved cerebellar representation produces more precise motor timing execution. On the other hand. activation of the visual cortex during generalization suggests an internal top-down recreation of the visual rhythmic pattern that facilitates execution of the tapping sequence.

The closest antecedent to the present study is an article by Bueti and collaborators (Bueti et al. 2012) where they found that the hemodynamic response in visual cortex was larger in the Post-than the Pre-training session of a visual interval discrimination task. These learning-based changes in the input sensory area were accompanied by an increase in the white-matter connectivity and gray-mater volume in the cerebellum, which are concordant with our findings. However, these authors found that the generalization from visual to auditory time discrimination was associated with the activation of the left inferior parietal cortex (Bueti et al. 2012), which we did not observe in our Post > Pre activation maps. This discrepancy can be due to the differences in experimental design between studies, with a similar visual interval discrimination learning task, but different generalization tasks. In fact, we used a time production task that involved rhythmic tapping production, which demands a strong sensorimotor coordination not present in an interval discrimination task (Merchant, Zarco & Prado 2008, Honing et al. 2018).

Besides the heterogeneity in the learning-generalization profile of our subjects learning transfer observed in Learners, all subjects benefited from the exposure and execution of the SCT on the two sessions. This practice effect is associated with a specific activation of the CTBGc across all participants independently of the learning and generalization performance in the IDT and SCT. A previous study found similar results, namely, changes in SMA and the basal ganglia linked to two days of training during a taping Synchronization task (Jantzen et al. 2002). Therefore, these results suggest that the core timing network is constantly refining its ability to internally time isochronous events and drive the tapping behavior and that practice is critical for activating this circuit and keeping the neural clock attuned and properly functioning.

## 5 Conclusions

We found large individual differences in the learning process associated with the intense training of an interval discrimination task, with a group of Learners showing an increase in timing precision after a week of practice, a group of Non-Learners with similar discrimination thresholds across training, and a group with no signs of learning but that showed a practice-based decrease in Temporal Variability between the two sessions of the generalization paradigm. Specific transfer of learning was observed in Learners as an increase in temporal precision when they accessed the representation of the trained interval to produce internally driven rhythmic movements. Importantly, the CTBGc showed activity associated with the practice of rhythmic tapping that is independent of the process of Learning and Generalization, suggesting pervasive plastic changes in this circuit to maintain the timing machinery calibrated.

## Acknowledgments

We thank Jennifer Coull, Víctor de Lafuente and Juan Fernández for their fruitful comments on the manuscript. We also thank Luis Prado, Raúl Paulín, Erick Pasaye, Juan Ortiz, Leopoldo González, Luis Aguilar, and Alejandro de León, for their technical assistance.

